# Repeat investigation during social preference behavior is suppressed in male mice with prefrontal cortex *cacna1c* (Ca_v_1.2)-deficiency through the dysregulation of neural dynamics

**DOI:** 10.1101/2023.06.24.546368

**Authors:** Jonathan Hackett, Viraj Nadkarni, Ronak S. Singh, Camille L. Carthy, Susan Antigua, Baila S. Hall, Anjali M. Rajadhyaksha

**Author notes:** Corresponding author: Anjali M. Rajadhyaksha, Department of Pediatrics, Pediatric Neurology, Weill Cornell Medicine, New York, NY 10065, USA.

## Abstract

Impairments in social behavior are observed in a range of neuropsychiatric disorders and several lines of evidence have demonstrated that dysfunction of the prefrontal cortex (PFC) plays a central role in social deficits. We have previously shown that loss of neuropsychiatric risk gene *Cacna1c* that codes for the Ca_v_1.2 isoform of L-type calcium channels (LTCCs) in the PFC result in impaired sociability as tested using the three-chamber social approach test. In this study we aimed to further characterize the nature of the social deficit associated with a reduction in PFC Ca_v_1.2 channels (Cav1.2^PFCKO^ mice) by testing male mice in a range of social and non-social tests while examining PFC neural activity using *in vivo* GCaMP6s fiber photometry. We found that during the first investigation of the social and non-social stimulus in the three-chamber test, both Ca_v_1.2^PFCKO^ male mice and Ca_v_1.2^PFCGFP^ controls spent significantly more time with the social stimulus compared to a non-social object. In contrast, during repeat investigations while Ca_v_1.2^PFCWT^ mice continued to spend more time with the social stimulus, Ca_v_1.2^PFCKO^ mice spent equal amount of time with both social and non-social stimuli. Neural activity recordings paralleled social behavior with increase in PFC population activity in Ca_v_1.2^PFCWT^ mice during first and repeat investigations, which was predictive of social preference behavior. In Ca_v_1.2^PFCKO^ mice, there was an increase in PFC activity during first social investigation but not during repeat investigations. These behavioral and neural differences were not observed during a reciprocal social interaction test nor during a forced alternation novelty test. To evaluate a potential deficit in reward-related processes, we tested mice in a three-chamber test wherein the social stimulus was replaced by food. Behavioral testing revealed that both Ca_v_1.2^PFCWT^ and Ca_v_1.2^PFCKO^ mice showed a preference for food over object with significantly greater preference during repeat investigation. Interestingly, there was no increase in PFC activity when Ca_v_1.2^PFCWT^ or Ca_v_1.2^PFCKO^ first investigated the food however activity significantly increased in Ca_v_1.2^PFCWT^ mice during repeat investigations of the food. This was not observed in Ca_v_1.2^PFCKO^ mice. In summary, a reduction in Ca_v_1.2 channels in the PFC suppresses the development of a sustained social preference in mice that is associated with lack of PFC neuronal population activity that may be related to deficits in social reward.

## Introduction

A lack of motivation to engage in social behavior is a feature of several neuropsychiatric diseases, including schizophrenia (SCZ) and autism spectrum disorder (ASD) [1, 2]. There is accumulating evidence from experiments in genetically modified mice that altered activity of Ca_v_1.2 L-type calcium channels (LTCCs) in the central nervous system can influence social cognition, impaired in these psychiatric disorders [3, 4]. Humans with single nucleotide polymorphisms in *CACNA1C*, the gene encoding the Ca_v_1.2 subunit of LTCCs, the most common isoform found in the brain [5], are more likely to exhibit low extraversion [6] and impaired facial emotional recognition [7], suggesting that Ca_v_1.2 channels may influence social interaction. In mice, both Ca_v_1.2 deficient mice [8] and knock-in gain of function Ca_v_1.2 Timothy Syndrome mice [9], as well as Ca_v_1.2 heterozygous knock-out rats [10–13] display significant social deficits, suggesting that altered calcium influx can bring about dysregulation of neuronal processing.

The prefrontal cortex (PFC) has long been implicated in various aspects of social behavior in humans and mice [14, 15]. This is consistent with impaired PFC function in *CACNA1C* neuropsychiatric risk SNP carriers and neuropsychiatric disorders that manifest with symptoms that encompass a range of social deficits, with affected individuals demonstrating reduced preference for social stimuli, impairments in reciprocal social interaction, abnormal social approach behavior and decreased social reward learning [1, 2, 16–18].

A study from our laboratory has shown that focal knockdown of Ca_v_1.2 channels in the adult PFC is sufficient to induce social impairment in the three-chamber social interaction test [8]. Electrophysiological evidence in PFC slices of Ca_v_1.2 deficient mice reveals a cellular E/I imbalance [8], a pathological mechanism linked to impaired social behavior [19, 20]. In the present study, we aimed to identify the nature of the social deficit associated with a reduction in expression of Ca_v_1.2 channels in the PFC by exposing male PFC *cacna1c* (Ca_v_1.2) deficient mice to a range of social and non-social tasks. To gain a greater understanding of the resulting alterations in neuronal population activity, we used GCaMP6s fiber photometry, an *in vivo* calcium imaging technique, to record neuronal dynamics in the PFC of Ca_v_1.2 deficient mice during behavior. By mapping synchronous bulk changes in calcium-driven fluorescence onto interactions with same-sex conspecifics or objects, fiber photometry revealed differences in activity patterns resulting from focal knockout of Ca_v_1.2 in the PFC.

## Methods

### Animals

Homozygous *cacna1c* floxed (cacna1c^fl/fl^; [21]) adult male mice and their wild-type littermates, on a C57BL/6J background, were used for all experiments. Mice were provided with food and water ad libitum and maintained on a 12-hour light/dark cycle (from 7 A.M. to 7 P.M.). All animals were group housed with two-three social partners. All procedures were conducted in accordance with the Weill Cornell Medicine Institutional Animal Care and Use Committee rules.

### Stereotaxic Surgery

Six to eight-week-old mice were anesthetized with a ketamine (100mg/ml) and xylazine (20mg/ml) cocktail and mounted in a stereotaxic surgical apparatus (David 40 Kopf Instruments, Tujunga, CA). AAV2/2-Cre-GFP or AAV2/2-GFP was sterotaxically injected into the right prelimbic cortex of adult *cacna1c*^fl/fl^ mice to induce focal knockdown as previously described [8]. Stereotaxic coordinates for the prefrontal cortex were anteroposterior (AP): +2.3 mm, mediolateral (MV): ±1.7 mm, dorsoventral (DV): −2.8 mm; angled 30° 3 toward the midline in the coronal plane, adopted from Paxinos and Franklin (citation). A 30-gauge Hamilton syringe was used to inject the virus at a rate of 0.1μl/min for a total volume of 0.8μl /hemisphere. Mice were then co-injected at the same coordinates in the left hemisphere, with 0.8μl of AAV2/2-Cre-GFP or AAV2/2-GFP and 0.2μl of AAV1-Syn-GCaMP6s-WPRE-SV40 (Penn Vector Core). An optical fiber was implanted at approximately 0.2mm dorsal of the injection site and fixed to the head with dental cement. Mice recovered in their home-cage for five weeks prior to behavioral testing.

### Green Fluorescent Protein (GFP) immunohistochemistry

GFP immunocytochemistry was used to confirm placement of surgical injections and placement of the optical fiber. Animals were anesthetized with Euthasol and perfused transcardially with 4% paraformaldehyde. Brains were dissected, post-fixed overnight in 4% PFA, and cryo-protected in 30% sucrose at 4°C for at least 72 hours. Brains were sectioned at a thickness of 50μm using a sliding microtome and sections were incubated in chicken anti-GFP (1:5000; Aves Lab Inc.) primary antibody for 24 hours at 4°C. The sections were rinsed in 0.1M phosphate-buffer (PB) and incubated with donkey anti-chicken Alexa Fluor 488 (1:500; Life Technology, Carlsbad, CA) antibody for 1 hour at room temperature. Sections were imaged using an epifluorescent microscope (Leica DM550B with Leica Application Suite Advanced Fluorescence 3.0.0 build 8134 software, Leica Microsystems, Wetzlar, Germany). Animals with improper bilateral injection or optical fiber placement were excluded from behavioral data analysis.

### Behavior testing

#### Three-chamber social approach

Social approach behavior was tested using the three-chamber social interaction test as previously published [8]. The test apparatus consisted of a white rectangular apparatus measuring 65cm x 40cm x 25cm that was divided into three equal-sized chambers, with an inverted pencil cup in each of the outer chambers. Removable barriers could be used to block off openings between chambers. Before trials, animals were connected to the fiber-optic patch-cord placed in the center chamber and removal of barriers initiated the start of a trial by allowing the animal free access to all chambers. Each mouse had a 5-minute habituation trial where each of the inverted cups was empty. Habituation was followed by a 5 minute “sociability” test where a novel age matched male C57BL/6J stranger mouse was placed under one cup and a novel object (a large Lego piece) in the cup in the opposing chamber. Social preference was measured as the time spent in a 7cm zone surrounding the cup containing the stranger mouse relative to time spent in the equivalent zone around the cup with the novel object. The time spent in each zone was further broken up into “bouts”, where a bout is defined as a continuous period spent in the investigation zone. Once the mouse left the zone for more than five seconds, it’s return to the zone was considered a new bout. The experiment was recorded using a video-camera mounted above the apparatus and the position of the experimental mouse was tracked using Ethovision software (Noldus, Wageningen, the Netherlands). Measurements were obtained via automated analyses using Ethovision combined with hand scoring of time spent per bout and interaction behavior (sniffing/clawing at cup). Interactions greater than one second in duration were included for analysis.

#### Reciprocal Interaction Social Assay

The test animal was connected to the fiber-optic patch-cord and placed in a new, empty cage with the lid removed and allowed to explore freely for 5 minutes. An age-matched male C57BL/6J stranger mouse was introduced and nose-nose (2 mice actively interacting head-on) and nose-tail (an active mouse pursuing a passive mouse) interactions were hand scored over 5 minutes. The experiment was recorded using a video-camera mounted above the apparatus and the position of the experimental mouse was tracked using Ethovision software.

#### Y-maze forced alternation social test

The Y-maze consisted of three equal sized arms with a 120° angle between each arm. In a 7-minute learning trial, mice were connected to a fiber-optic patch-cord, placed in one arm and allowed access to one other open arm with a novel age-matched male C57BL/6J stranger mouse confined in a transparent glass cylinder. In the subsequent 7-minute test trial, the previously blocked arm containing a novel object confined in an identical glass cylinder was opened and the mouse could explore all three arms. The time spent in each arm was the metric used to measure tendency to explore novel environments. The experiment was recorded using a video-camera mounted above the apparatus and the position of the experimental mouse was tracked using Ethovision software.

#### Linear track social approach

The three-chamber apparatus was adapted into a 70cm x 12cm x 25cm linear track, with a novel age-matched male C57BL/6J stranger mouse under an inverted pencil cup on one end of the track and a novel object under a cup on the other end. The experimental protocol is identical to the three-chamber task, with a 5-minute habituation phase followed by a 5-minute test phase. The behavior recorded were the transitions from the object and the stranger mouse and vice versa to measure approach behavior [22]. Transitions greater than 6 seconds and less than 2 seconds were excluded from analysis (<25% data) to ensure comparison across a similar time frame. The experiment was recorded using a video-camera mounted above the apparatus and the position of the experimental mouse was tracked using Ethovision software. Measurements were obtained via automated analyses using Ethovision combined with hand scoring of time spent per bout and interaction behavior (sniffing/mounting cup).

#### Modified three-chamber food reward approach

The three-chamber social interaction test was modified to test a nonsocial stimulus. The protocol described above was repeated where the novel stranger mouse was replaced with a novel object that had been allowed to soak in wet food mesh for 10 minutes. Preference for the food-smelling novel object was measured as the time spent in a 2.5” zone surrounding the cup relative to time spent in the equivalent zone around the cup with the neutral-smelling novel object.

### Fiber photometry and analysis

Fiber photometry was performed as described previously (citation). For analysis, raw data was processed using custom MATLAB software, which normalized the fluorescent signal by calculating ΔF/F from a baseline (f) of the median of the data points in a 30s sliding time window. To account for variability in the dynamic range of the signal across animals and trials, ΔF/F values were converted into a z score calculated using the median across the trial:

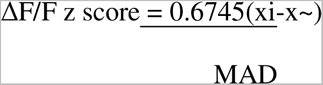

MAD denotes the median absolute deviation, xi the ΔF/F value and x~ the median. Neural activity was typically quantified as the summed z-score ΔF/F over a significance level greater than 1 standard deviation divided by the sample time. Animals with a signal that was sufficiently depleted (maximum ΔF/F < 5%) were excluded from analysis.

### Statistical analysis

Behavioral and fiber photometry group data are presented as mean with individual data points or ± SEM. ANOVA and post hoc Bonferroni tests were used to analyze across stimuli. Student’s t-tests were used for two groups and one-sample t-tests to analyze transitions in the linear track task. GraphPad Prism (La Jolla, Ca, USA) was used for all statistical analyses. Tableau Desktop 2018.2 (Seattle, WA, USA) was used for data visualization.

## Results

### Focal knockout of Ca_v_1.2 channels in the prefrontal cortex prevents mice from developing a social preference during investigation behavior

We have previously demonstrated that mice with a deficiency of *cacna1c* (Ca_v_1.2) in the prefrontal cortex (Ca_v_1.2^PFCKO^) have impaired sociability in the three-chamber social interaction test [8], measured as time spent in a social ‘investigation zone’ extending 2.5” around the inverted cup containing the conspecific relative to the equivalent object zone (**Figure 1A**). To further elucidate the specific nature of the social deficit, we focused on how social preference changed over time during the 5-minute testing period. To this end, homozygous *cacna1c* floxed (*cacna1c*^fl/fl^; [21]) adult male mice were bilaterally stereotaxically injected with AAV2/2-Cre-GFP into the PFC to induce focal knockout (Ca_v_1.2^PFCKO^) (**Figure 1B, left**). A second cohort of mice were injected with AAV2/2-GFP to generate Ca_v_1.2^PFCGFP^ control mice. Additionally, to query prefrontal neuronal activity during social behavior both cohorts were injected unilaterally with the genetically encoded calcium indicator GCaMP6s driven by the synapsin promoter (AAV1-Syn-GCaMP6s-WPRE-SV40) (**Figure 1B, right**). By implanting an optic-fiber above the injection site we were able to record bulk changes in fluorescence in the soma of neurons during behavior.

**Figure 1.**
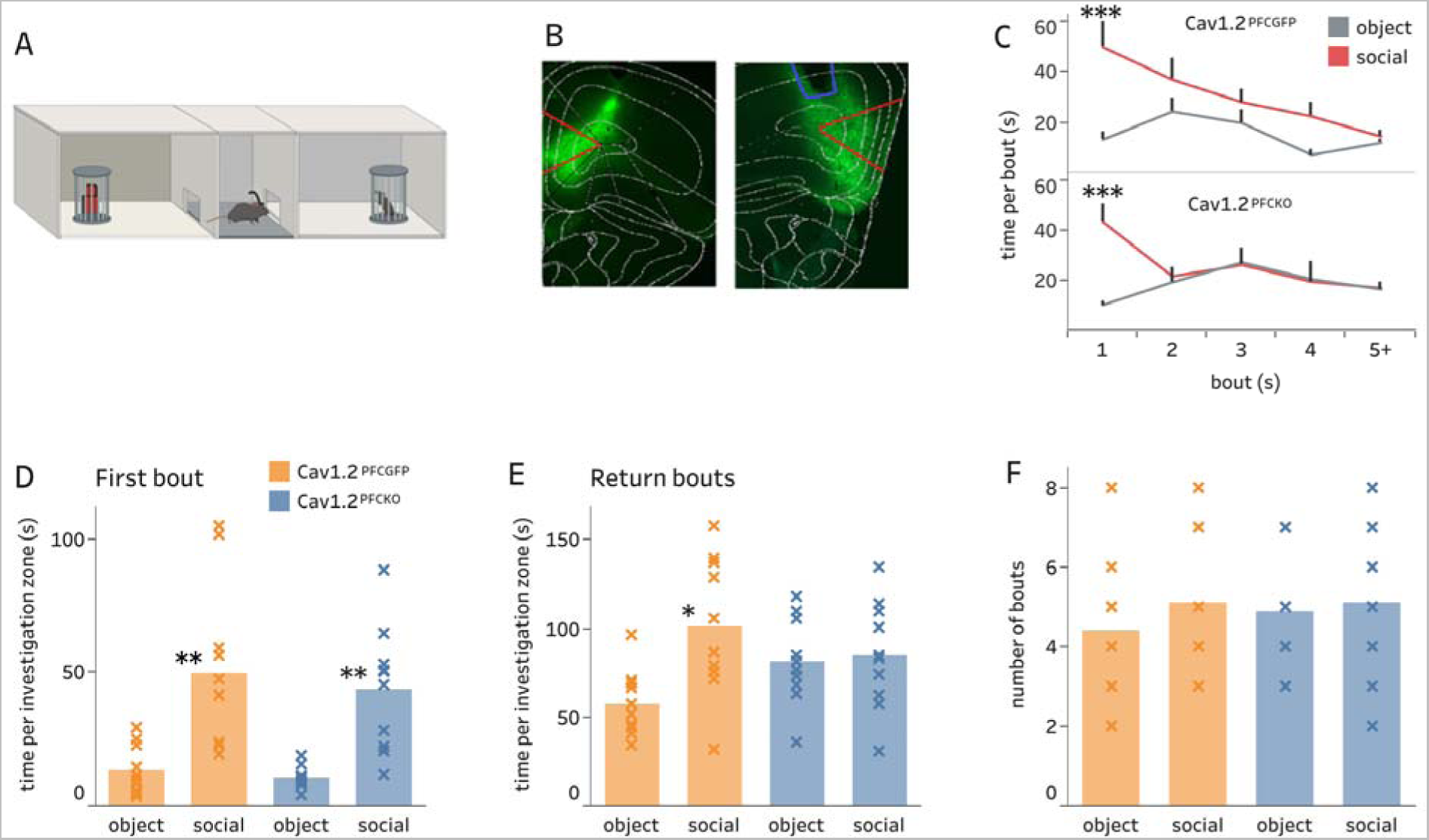
Reduction of Ca_v_1.2 in the prefrontal cortex generates a deficit in sociability after the first encounter with a conspecific. **1A.** Three-chamber social approach apparatus. **1B.** Exemplar image of GFP expression at injection site, taken at Bregma +2.3mm with prelimbic cortex outlined in red for reference. Left: AAV-Cre-GFP expression of virus. Right: AAV-Cre-GFP and GCaMP6s expression with optic fiber placement highlighted in blue. **1C.** The average time per bout for the novel object and social partner. Ca_v_1.2^PFCGFP^ mice spend significantly more time with the social stimulus than the object on first bout (Bonferroni *post hoc* test, ***p = 0.0002) and continue to spend more time with the social partner on subsequent bouts. In contrast Ca_v_1.2^PFCKO^ mice spend significantly more time with social than object on first encounter (Bonferroni *post hoc* test, ***p = 0.0002) but show no preference between the two stimuli in subsequent bouts (Ca_v_1.2^PFCGFP^ n=10, Ca_v_1.2^PFCKO^ n=10). **1D.** Social preference during first investigation bout. Both animal groups show a strong social preference (Bonferroni *post hoc* test, Ca_v_1.2^PFCGFP^: **P=0.0017 vs object, n=10; Ca_v_1.2^PFCKO^: **p=0.0050 vs. object, n=10). **1E.** Social preference during return bouts when revisiting stimuli. Ca_v_1.2^PFCGFP^ mice retained a social preference (Bonferroni *post hoc* test, Ca_v_1.2^PFCGFP^: *p=0.0119 vs object, n=10), whereas Ca_v_1.2^PFCKO^ mice exhibited a deficit in sociability (p > 0.9999, n=10). **1F.** The number of visits to each zone or ‘bouts’ was not significantly different between groups (Ca_v_1.2^PFCGFP^ n=10, Ca_v_1.2^PFCKO^ n=10). Error bars ± s.e.m.

Three weeks following injection, to measure sociability in these mice, animals were tested in the three-chamber social approach test containing a novel mouse in one end of the chamber and an object in the opposite end of the chamber (**Figure 1A**). Social and object investigations were divided into bouts, defined as a continuous period of time spent in either the social or object investigation zone upon entry into that zone, over the 5-minute testing period. Examining the time spent per investigation bout over the 5 minutes revealed that although Ca_v_1.2^PFCGFP^ mice habituated to the social stimulus over time, they continued to spend more time with the social stimulus compared to the object, with significantly more time during the first bout (**Figure 1C**, top; *two-way* ANOVA, main effect of time per bout F_4,71_=3.007, p =0.0237; main effect of zone F_1,71_=12.14, p=0.0009). In contrast, the difference in time spent with the social stimulus compared with the object in Ca_v_1.2^PFCKO^ mice was indistinguishable after the initial social encounter (**Figure 1C bottom**; *two-way* ANOVA, significant interaction F_4,74_ = 4.092, p=0.0047, no main effect of time per bout F_4,74_=1.608, p =0. 1813; main effect of zone F_1,74_=4.917, p=0.0297).

Next to further define aspects of social approach behavior that may be disrupted in Ca_v_1.2^PFCKO^ mice, we divided the test into the first bout where investigation is exploratory in a novel environment and return bouts where the stimuli is no longer novel. During the first bout, both Ca_v_1.2^PFCGFP^ control and Ca_v_1.2^PFCKO^ mice spent significantly more time with the novel social conspecific relative to the novel object (**Figure 1D**; *two-way* ANOVA, main effect of zone, F_1,36_ = 29.44, p<0.0001; no main effect of PFC KO, F_1,36_ = 0.5443, p=0.4655). For return bouts, Ca_v_1.2^PFCGFP^ control mice continued to spend significantly more time in the social investigation zone, whereas Ca_v_1.2^PFCKO^ mice spent similar time with the social stimulus as with the novel object **(Figure 1E**; *two-way* ANOVA, significant interaction F_1,36_ = 4.704, p=0.0368). This genotype difference was not a result of difference in number of bouts as there was no difference in bouts into the object or social zone between Ca_v_1.2^PFCGFP^ and Ca_v_1.2^PFCKO^ mice (**Figure 1F**). These results demonstrated that PFC Ca_v_1.2 deficiency does not alter initial exploration, nor number of investigations but does result in decrease in time spent during repeat investigations of the social stimulus, suggesting a more specific role for PFC Ca_v_1.2 channels in social interaction behavior.

### Focal knockdown in **Ca_v_**1.2 channels in the PFC is concurrent with reduced neuronal population activity during repeat social interactions

To determine whether the observed deficit in revisiting social stimuli in the three-chamber test is reflected in altered prefrontal neuronal activity, we employed *in vivo* calcium recordings via fiber photometry [23, 24] to acquire real-time population activity of PFC neurons during social behavior. Synchronization of the fluorescent signal with the movements of the mice allowed us to timestamp the real-time dynamics of PFC neurons onto different features of social behavior: investigation of the social or object zone and interactions while in the zone (e.g., sniffing/clawing) (**Figure 2A**). Due to variations in the dynamic range of the fluorescent signal, the ΔF/F was z-scored using the median across the trial. Neural activity was quantified as the summation of z-scored ΔF/F over a significance level greater than 1 standard deviation, divided by the sample time (**Figure 2B**). It is important to note that there was no difference in the average GCaMP6s fluorescence of the PFC neurons between the Ca_v_1.2^PFCGFP^ and Ca_v_1.2^PFCKO^ mice during the entire habituation period (**Supplementary Figure 1**) or the 5-minute three-chamber test (data not shown) indicating that loss of Ca_v_1.2 Ca^2+^channels does not affect the bulk GCaMP6s signal.

**Figure 2.**
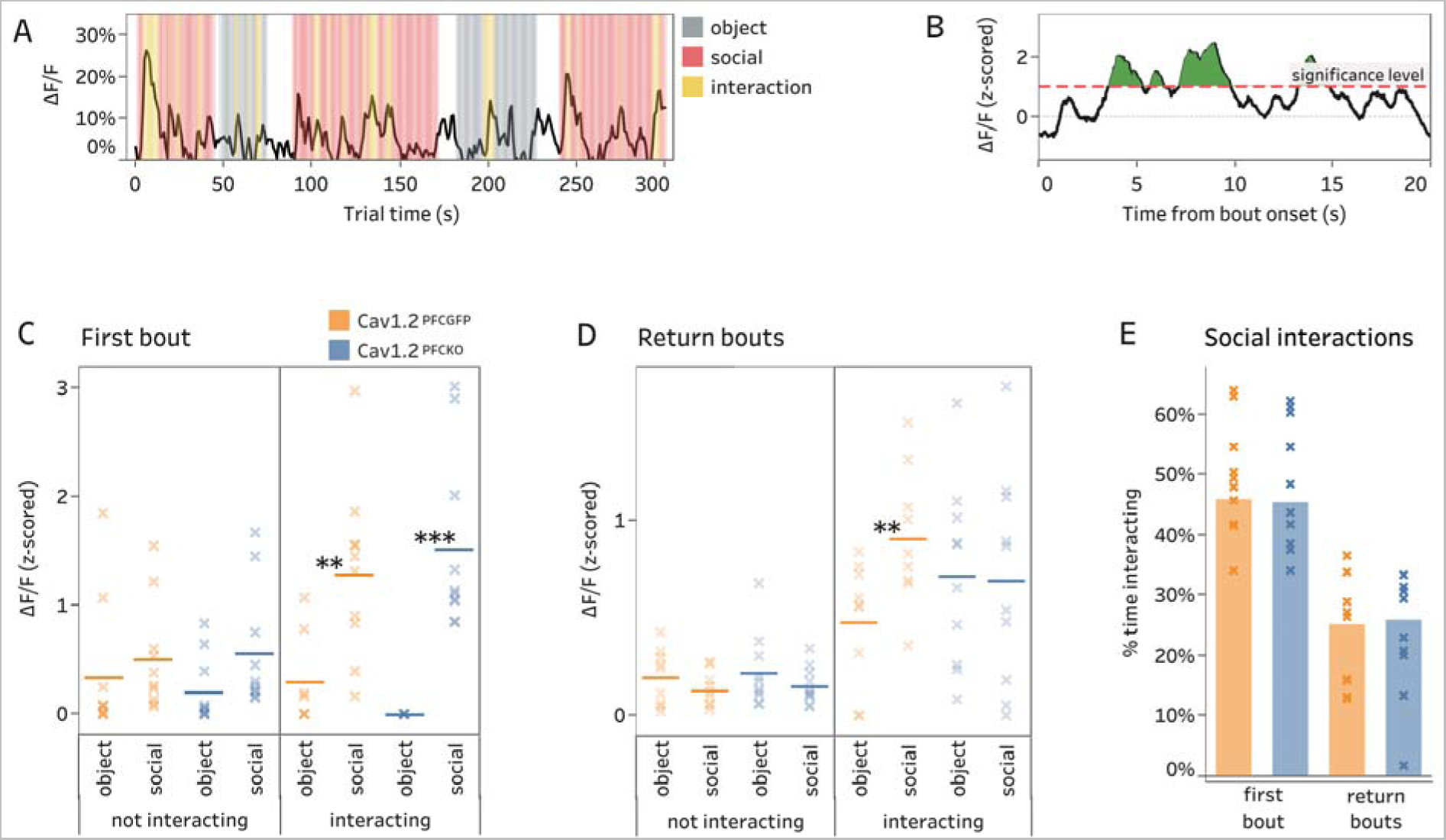
Ca_v_1.2 PFC knockout mice show deficit in neural activity during social interactions upon repeat investigation. **2A.** Exemplar fiber photometry ΔF/F trace for a 5-minute three chamber trial. Hand scored behavior is overlayed: social bouts (red), object bouts (grey) and within-bout interactions with the mouse or object (yellow). **2B.** Schematic of quantification of fiber photometry signal. Raw ΔF/F was converted into a Z-score and activity defined as the average greater than one standard deviation. **2C.** Quantification of ΔF/F during the first bout, while mice are not interacting and interacting with the stimulus (object or social). Right, Interacting. ΔF/F was greater in the PFC of Ca_v_1.2^PFCGFP^ and Ca_v_1.2^PFCKO^ while interacting with the stimulus during the first social bout relative to the first object bout (Bonferroni *post hoc* test, Ca_v_1.2^PFCGFP^: **p=0.0038 vs. object; n=10, Ca_v_1.2^PFCKO^: ***p=0.0006 vs. object, n=10). Left, not interacting. No difference in signal was seen when mice were not interacting for either genotype. **2D.** Quantification of ΔF/F during interactions in return bouts. Right, Interacting. ΔF/F was significantly greater during social interactions relative to object interactions in Ca_v_1.2^PFCGFP^ mice with no difference in Ca_v_1.2^PFCKO^ mice (linear mixed effects model, **p=0.00478, number of bouts= 47, n= 6; Ca_v_1.2^PFCKO^: p=0.7398 number of bouts=66, n= 8). Left, Not interacting. No significant difference in ΔF/F when Ca_v_1.2^PFCGFP^ or Ca_v_1.2^PFCKO^ mice were not interacting. **2E.** Percentage of time spent interacting with social partner for first bout and return bouts. Ca_v_1.2^PFCGFP^ and Ca_v_1.2^PFCKO^ mice spent similar time interacting with the novel social partner during first and return bouts.

During the first bout of the three-chamber test, the GCaMP6s fluorescent signal recorded in the PFC during active investigation of the social stimulus was significantly higher than that recorded during active object investigation irrespective of viral treatment (**Figure 2C, interacting,** *two-way* ANOVA, main effect of zone, F_1,28_ = 35.24, p<0.0001; no main effect of zone, F_1,28_ = 0.3312, p=0.5696). No difference in PFC activity was observed when mice were not actively investigating while in the contact zone (**Figure 2C, not interacting**). In Ca_v_1.2^PFCGFP^ control mice, the elevated PFC neuronal activity during active social investigations relative to object investigations was maintained in return bouts (**Figure 2D, interacting).** In contrast, in Ca_v_1.2^PFCKO^ mice the higher neuronal activity between social and object during return bout active investigations was no longer observed (**Figure 2D, interacting**). No difference was seen for either viral group when mice were not interacting (**Figure 2D**). The lack of an increase in PFC neural activity in Ca_v_1.2^PFCKO^ mice was not a result of lower time spent interacting with the social stimulus compared to Ca_v_1.2^PFCGFP^ mice (**Figure 2E**). These results demonstrated that loss of Ca_v_1.2 channels in the PFC results in lack of an increase in PFC neuronal activity time locked to active social investigations during return bouts.

### Prefrontal cortex activity upon repeat social investigations is predictive of social preference behavior in control mice but not **Ca_v_**1.2^PFCKO^ mice

To more comprehensively link the deficit in social investigation in Ca_v_1.2^PFCKO^ mice with the altered neuronal dynamics, we evaluated whether the fiber photometry signal upon entry into investigation zone was predictive of place preference, measured as the subsequent time spent in the zone. We defined the ‘entry’ period as the time 1 second before and 3 seconds after entering the investigation zone (**Figure 3A**), thereby encompassing PFC activity during the transition into the zone and the initial interaction.

**Figure 3.**
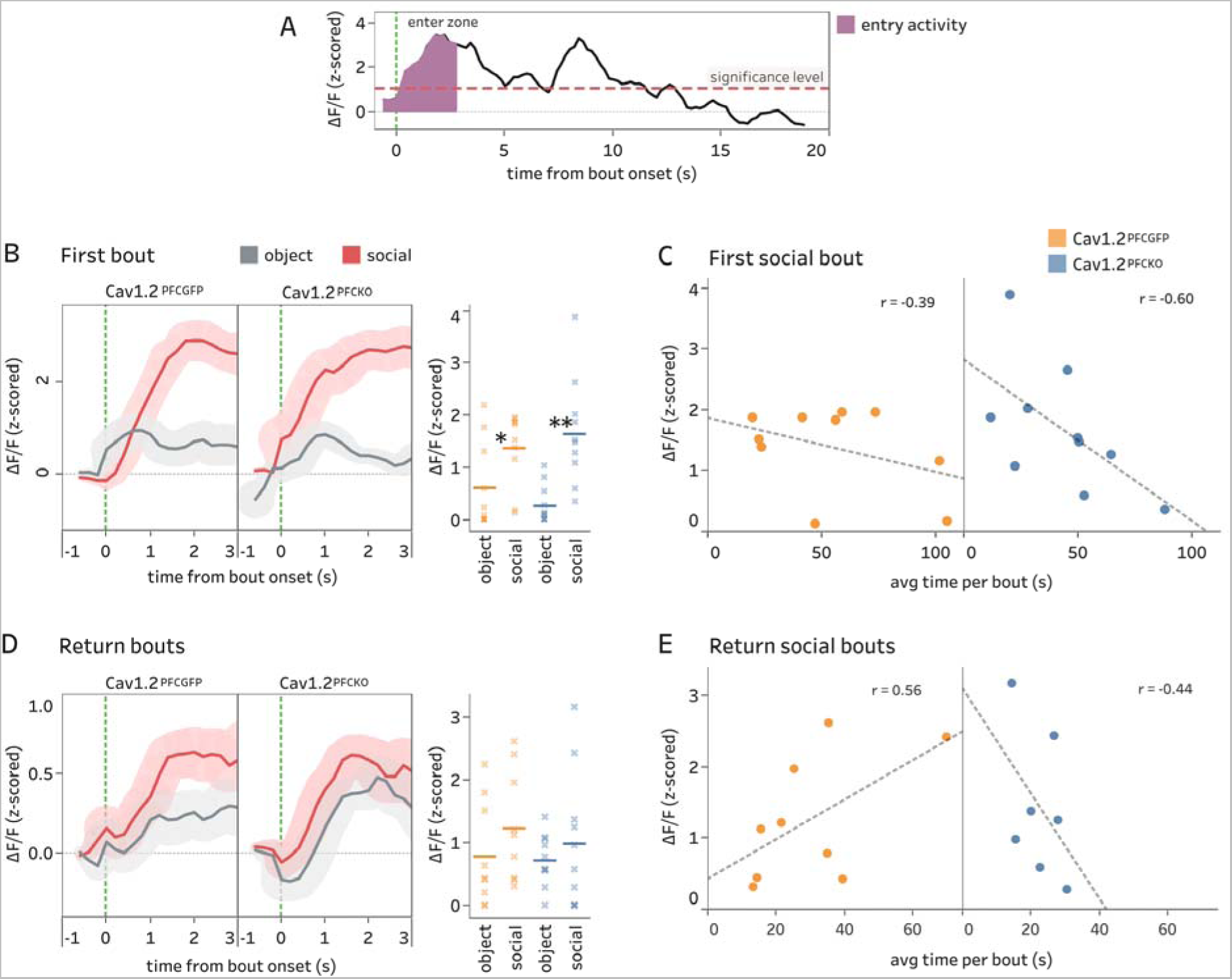
Prefrontal activity correlates with time spent revisiting social stimuli in Ca_v_1.2^PFCGFP^ mice and time spent during first investigation in Ca_v_1.2^PFCKO^ mice. **3A.** Example fiber photometry ΔF/F trace for entrance into an investigation zone. ‘Entry’ signal (purple) quantified as average ΔF/F above one standard deviation from 1 second before entering zone to 3 seconds after. **3B.** Left: average ΔF/F traces at the point of entering the investigation zone during the first bout (social – red, object - grey). Shaded region ± s.e.m. Right: quantification of entry signal (−1s to +3s from entering zone). Both groups showed an elevated ΔF/F signal on entering the zone but with no significant difference across viral treatment (Bonferroni *post hoc* test, *p=0.0252, **p=0.0018; Ca_v_1.2^PFCGFP^ n=10, Ca_v_1.2^PFCKO^ n=10). **3C.** Correlation plot of entry GCaMP6s signal in first social bout with time spent in the investigation zone for that same bout. The social entry ΔF/F in Ca_v_1.2^PFCKO^ mice is negatively correlated with the subsequent time spent in the investigation zone, whereas there is no significant relationship in Ca_v_1.2^PFCGFP^ mice (linear regression, *P<0.05; Ca_v_1.2^PFCGFP^ n=10, Ca_v_1.2^PFCKO^ n=10). **3D.** Left: average ΔF/F traces at the point of entering the investigation zone during return bouts (social - red, object - grey). Shaded region ± s.e.m. Right: quantification of signal (−1s to +3s from entering zone). ΔF/F was higher in Ca_v_1.2^PFCGFP^ mice but did not reach significance, linear mixed effects model, p=0.4529, number of entries = 61, n=10). There was an equal rise in fluorescence in Ca_v_1.2^PFCKO^ mice as they approached both the social and object with no significant difference between the two (linear mixed effects model, p=0.4973, number of entries=80, n=10). **3E.** Correlation plot of entry signal in social return bouts with time spent in the investigation zone for that same bout. The social entry ΔF/F in Ca_v_1.2^PFCGFP^ mice was positively correlated with the subsequent time spent in the investigation zone, whereas there is no significant relationship in Ca_v_1.2^PFCKO^ mice (linear regression, **P<0.01; Ca_v_1.2^PFCGFP^ n=18, Ca_v_1.2^PFCKO^ n=16). Bouts that recorded ΔF/F over significance level (1 standard deviation) included.

Plotting average ΔF/F as mice enter the zone during the first bout demonstrated that the increase in activity during the first social investigation was significantly greater than the response to the object in both Ca_v_1.2^PFCGFP^ and Ca_v_1.2^PFCKO^ animals (**Figure 3B**; *two-way* ANOVA, main effect of zone, F_1,36_ = 19.22, p<0.0001, no main effect of PFC KO, F_1,36_ = 0.003718, p=0.9517). Yet during this first social investigation of a novel conspecific, there was no correlation in entry activity with the time spent with the stranger mouse in the Ca_v_1.2^PFCGFP^ mice, while the entry PFC activity in Ca_v_1.2PFC^KO^ mice was negatively correlated to the time spent with the stranger mouse (**Figure 3C**). We additionally looked at the entry signal in return bouts of the three-chamber social approach test (**Figure 3D**). For Ca_v_1.2^PFCGFP^ mice, when comparing the PFC activity when entering the social zone versus the object zone, the PFC activity was greater upon entering the social zone compared to object zone (**Figure 3D, left**), however the average magnitude of the increase did not reach significance (**Figure 3D, right**). For Ca_v_1.2^PFCKO^ mice there was an increase in PFC activity upon entry into the social and the object zone (**Figure 3D, left**) with no difference in average magnitude (**Fig 3D, right**). However, interestingly in Ca_v_1.2^PFCWT^ mice there was a positive correlation between the social entry ΔF/F signal and the amount of time spent in the subsequent bout in control mice (**Figure 3E, left**). There was no correlation found in Ca_v_1.2^PFCKO^ mice (**Figure 3E, right**).

### Focal knockout of PFC **Ca_v_**1.2 does not generate a deficit in reciprocal social interactions with a free-roaming mouse or prevent a preference for novel environments in a forced alternation Y-maze test

Next to further define the role of the PFC in social interactions and the impact of Ca_v_1.2 deficiency therein we first assessed reciprocal social interactions with a free-roaming mouse (**Figure 4A**). Subject mice were habituated to a novel cage for 5 minutes before the introduction of a novel conspecific. Nose-tail (pursuit and receiving) and nose-nose (sniffing) interactions were scored to measure sociability. In contrast to the three-chamber test, Ca_v_1.2^PFCKO^ mice had similar number of interactions for all three measures as that to Ca_v_1.2^PFCGFP^ control mice, demonstrating similar sociability (**Figure 4B**). Examination of PFC activity using fiber photometry time-locked to each of the social interaction measures found no difference in activity between groups during the first interaction (**Figure 4C)** or any subsequent interactions (**Figure 4D**).

**Figure 4.**
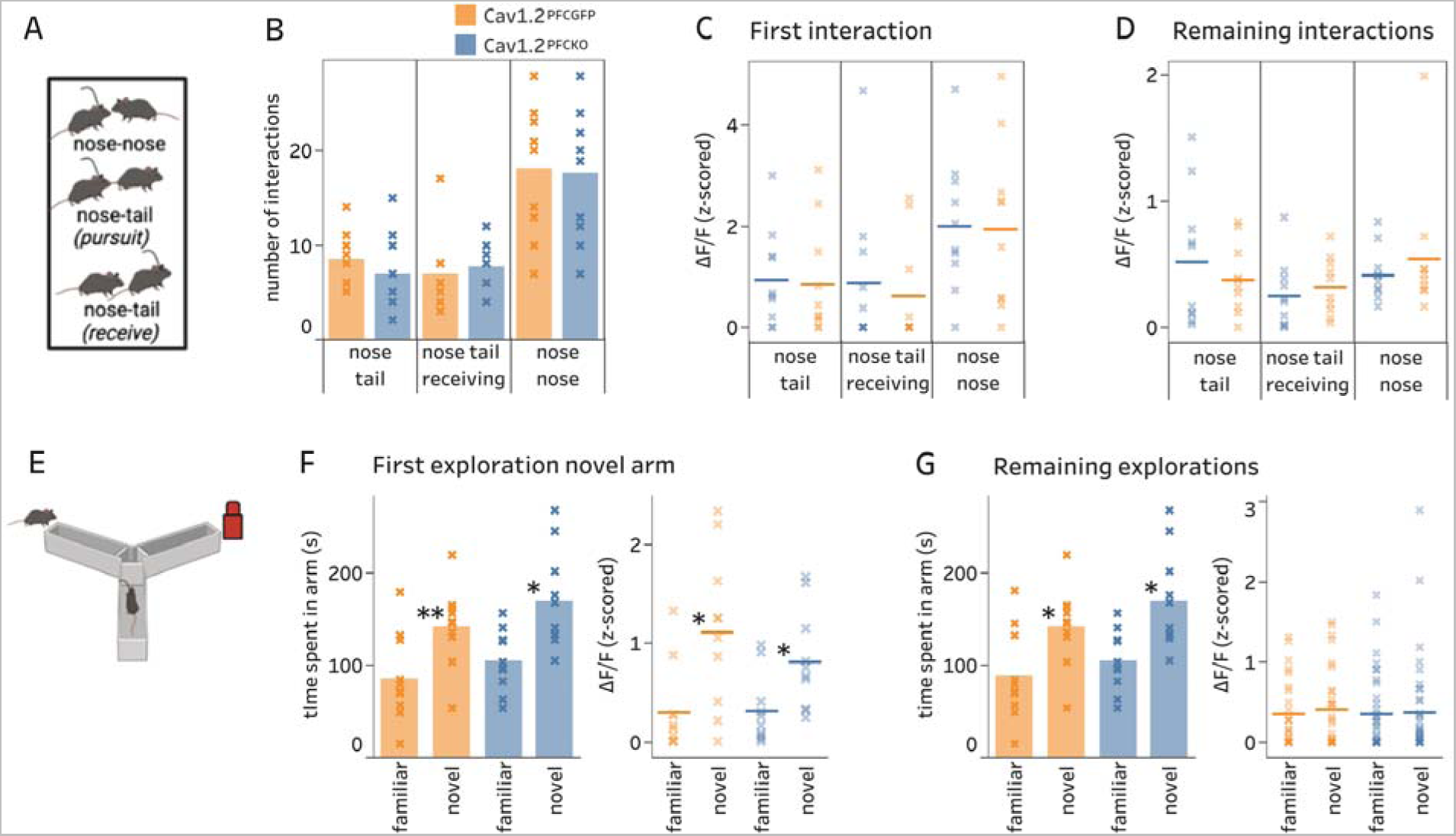
Reciprocal social interaction assay and Y-maze forced alternation test produced no behavioral deficits or discrepancies in neural dynamics in Ca_v_1.2^PFCKO^ mice. **4A.** Reciprocal social interaction assay apparatus. **4B.** Nose-tail pursuit, nose-tail receiving and nose-nose, interactions were recorded between a subject mouse and the stranger mouse, freely moving in a cage to test reciprocal interactions. There were no statistical differences in social interactions between Ca_v_1.2^PFCGFP^ and Ca_v_1.2^PFCKO^ mice. **4C.** ΔF/F during first social interactions, with no differences between Ca_v_1.2^PFCKO^ and Ca_v_1.2^PFCGFP^ mice. **4D.** ΔF/F during any remaining social interactions after the first, with no differences between KO and Ca_v_1.2^PFCGFP^ mice. **4E.** Y-maze forced alternation apparatus. **4F.** First exploration of novel object arm. Left, Both Ca_v_1.2^PFCGFP^ and Ca_v_1.2^PFCKO^ mice showed a preference for exploring the novel object arm on first exploration (Bonferroni *post hoc* test, Ca_v_1.2^PFCGFP^: **p=0.0064 and Ca_v_1.2^PFCKO^: *p=0.0216 vs. familiar social, n=10). Right, Ca_v_1.2^PFCGFP^ and Ca_v_1.2^PFCKO^ mice had significantly higher signal in the novel object arm compared to familiar social arm (Bonferroni *post hoc* test, Ca_v_1.2^PFCGFP^: *p=0.0130 and Ca_v_1.2^PFCKO^: *p=0.0146 vs. familiar social, n=10). **4G.** Return explorations of novel object arm. Left, Both Ca_v_1.2^PFCGFP^ and Ca_v_1.2^PFCKO^ mice continue to show a preference for exploring the novel object arm over the course of the 7-minute trial (Bonferroni *post hoc* test, Ca_v_1.2^PFCGFP^: *p=0.0425 and Ca_v_1.2^PFCKO^: *p=0.0162 vs. familiar social, n=10). Right, There was no associated increase in ΔF/F after the first exploration in either genotype (Linear mixed effects model, Ca_v_1.2^PFCGFP^: p = 0.5902, number of explorations = 52, n = 10; Cav1.2^PFCKO^: p = 0.9048, number of explorations = 68, n = 10).

Next, we tested mice in a modified Y-maze forced alternation test (**Figure 4E**) that tests working memory and recruits the PFC [25, 26]. Mice were given seven minutes to explore one arm containing a conspecific on a slightly raised glass cylinder that permitted no interaction. For the test phase, mice were additionally permitted access to a novel arm containing an object in an equivalent position (**Figure 4E**). Time spent exploring the novel object arm was measured. Ca_v_1.2^PFCGFP^ and Ca_v_1.2^PFCKO^ mice spent significantly more time in the novel object arm relative to the familiar social arm on first exploration (**Figure 4F, left.** *two-way* ANOVA, main effect of zone, F_1,36_ = 22.28, p<0.0001, no main effect of PFC KO, F_1,36_ = 0.4862, p=0.4901), which was mirrored by an elevated PFC activity during time spent with object in the novel arm (**Figure 4F, right.** *two-way* ANOVA, main effect of zone, F_1,36_ = 14.22, p=0.0006, no main effect of PFC KO, F_1,36_ = 0.7033, p=0.4072). Preference for the novel arm was sustained in return investigations for both groups (**Figure 4G, left,** *two-way* ANOVA, main effect of zone, F_1,36_ = 17.20, p=0.0002, no main effect of PFC KO, F_1,36_ = 2.373, p=0.1322), although PFC activity was no longer higher during novel arm exploration (**Figure 4G, right**).

### **Ca_v_**1.2 PFC channel deficiency leads to disrupted spatially modulated firing of PFC neuronal populations during social approach behavior

To focus specifically on approach behavior, we tested mice in a modified version of the three-chamber test, where animals were placed on a 70cm long, 12cm wide linear track, with a novel social stimulus confined in a cage at one end and a novel object stimulus at the opposite end (**Figure 5A**). This apparatus allowed us to record the fiber photometry signal relative to mouse’s position on the track and because the animal always had a clear sight of the target (not hidden behind a wall), any movement in that direction was considered approach behavior towards either the object or social cages. These movements from one end of the linear track to the other were scored as a ‘transition to object’ or ‘transition to social’ (**Figure 5B**).

**Figure 5.**
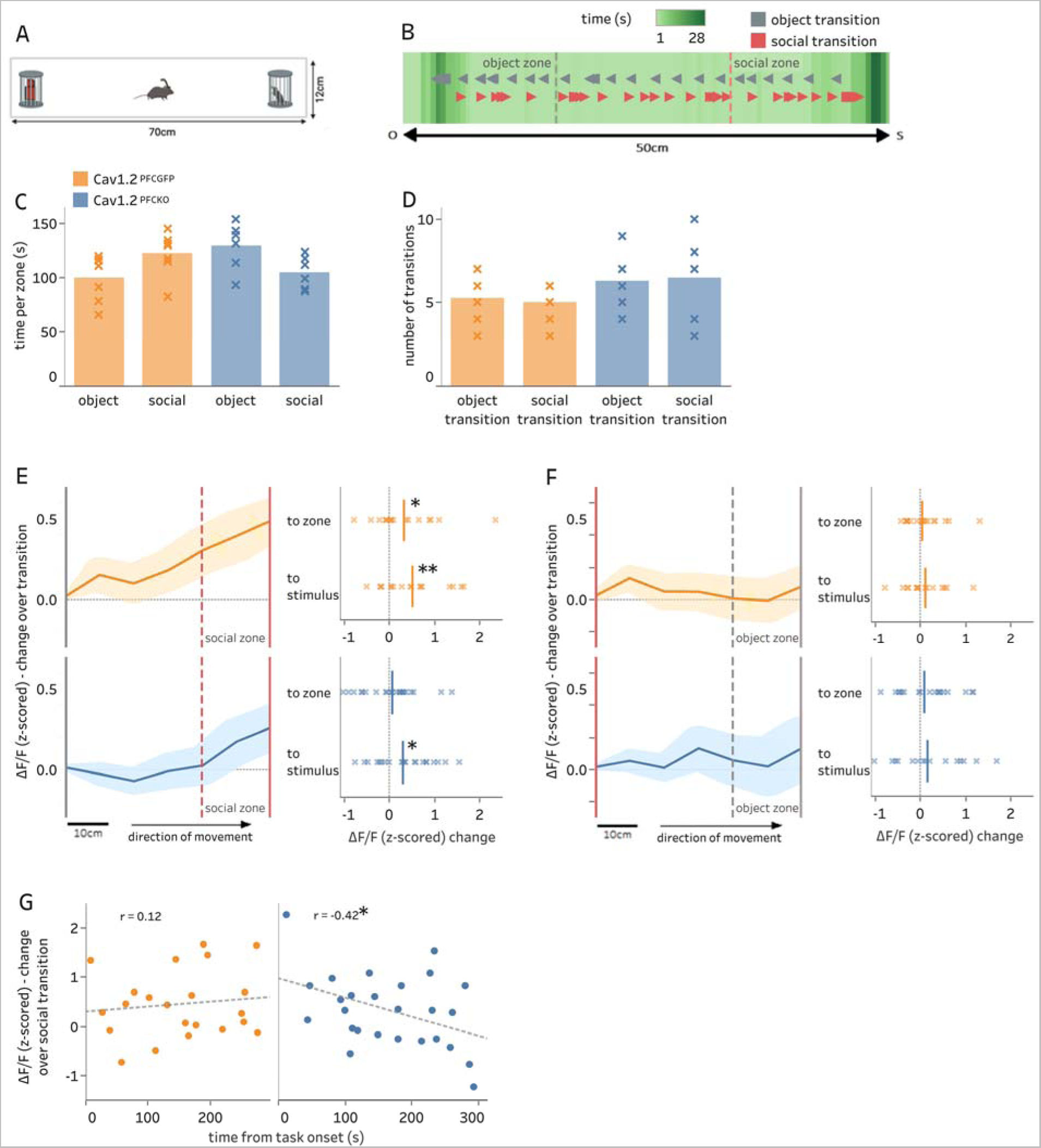
PFC activity in the linear track social approach test is delayed in PFC Ca_v_1.2 knockout mice. **y** Linear track social approach apparatus. **5B.** Heat map of time spent in each part of the linear track. Transitions between the novel object and stranger mouse (examples overlaid) were tracked and recorded to quantify the change in neural dynamics during social approach. **5C.** Time spent in social and object zones at the end of the track. Ca_v_1.2^PFCGFP^ mice displayed higher preference for the social stimulus whereas Ca_v_1.2^PFCKO^ mice displayed a diminished social preference (Ca_v_1.2^PFCGFP^ n=7, Ca_v_1.2^PFCKO^ n=6). **5D.** The number of transitions to the object or social zone was not significantly different between groups (Ca_v_1.2^PFCGFP^ n=7, Ca_v_1.2^PFCKO^ n=6). **5E. Left,** Average traces of the change in ΔF/F from the point of leaving the novel social stimulus (stranger mouse) to reaching the object zone during a transition. Transitions to object with quantified change in ΔF/F up to edge of object zone and to the object. Shaded region ±s.e.m. First encounters with stimuli are excluded. In Ca_v_1.2^PFCGFP^ and Ca_v_1.2^PFCKO^ mice, the increase in the fiber photometry signal was significantly greater than zero only during social approach stimulus (one-sample t-test, *P<0.05, **P<0.01; Ca_v_1.2^PFCGFP^ n=17, Ca_v_1.2^PFCKO^ n=22). In Ca_v_1.2^PFCGFP^ mice, the increase in signal preceded that of Ca_v_1.2^PFCKO^ mice in being significantly greater than zero at the point of reaching the social zone boundary located 14cm from the stimulus (one-sample t-test, *P<0.05; n=17). **5F.** Correlation plot of increase in ΔF/F during transitions to social with time point over the course of a five-minute trial. Ca_v_1.2^PFCKO^ mice showed a negative correlation (linear regression, R-squared = 0.17, *P<0.05), indicating a decrease in prelimbic ΔF/F during transitions to social over time.

First, we measured if the social behavior observed in the three-chamber test was reproducible in the linear track. The time spent with the social stimulus in the zone at one end of the track was higher compared to the object zone at the opposite end of the track in Ca_v_1.2^PFCGFP^ mice but not Ca_v_1.2^PFCKO^ (**Figure 5C**; *two-way* ANOVA, significant interaction (genotype x zone) F_1,22_ = 8.862, p=0.0070) consistent with the social deficit seen in the three chamber social test in Ca_v_1.2^PFCKO^ mice [8]. There was no difference in the number of transitions to either the social or object zone for either animal group (**Figure 5D**). We next examined the change in fluorescence as mice transitioned from the object zone (**Figure 5E** left, grey solid line) to the social zone (**Figure 5E** left, red dashed line) and the social stimulus (**Figure 5E** left, red sold line), as well as transitions from the social zone (**Figure 5F** left, red solid line) to the object zone (**Figure 5F left**, grey dashed line) and object stimulus (**Figure 5F left,** grey solid line). Looking at transitions to the social zone (**Figure 5E**), we observed a significant change in approach ΔF/F signal when Ca_v_1.2^PFCGFP^ mice transitioned to the social zone as well as the social stimulus (**Figure 5E**, right). The signal reached significance at the point of the social zone boundary (located 14cm from the conspecific). In contrast Ca_v_1.2^PFCKO^ mice did not demonstrate a significant change in approach ΔF/F signal when they approached the social zone but there was a change when they approached the social stimulus (**Figure 5E**, right). Thus, in Ca_v_1.2^PFCGFP^ mice the ramp-up in signal during social approach preceded that seen in Ca_v_1.2^PFCKO^ mice. Also, as compared to the ΔF/F signal seen in Ca_v_1.2^PFCGFP^ mice, the approach signal declined over the course of the trial in Ca_v_1.2^PFCKO^ mice as they approached the social zone (**Figure 5G**). There was no change in ΔF/F signal when both Ca_v_1.2^PFCGFP^ and Ca_v_1.2^PFCKO^ mice approached the object zone or object stimulus (**Figure 5F**). Taken together, a reduction of Ca_v_1.2 channels in the prefrontal cortex delays the increase in PFC approach activity when revisiting social stimuli and furthermore PFC activity declines over time during transitions to social.

### **Ca_v_**1.2 channel deficiency does not disrupt non-social food reward but results in PFC activity deficit

The behavioral deficit established in the three-chamber social approach test is feasibly related to an impairment in reward processing [14]. To determine whether the altered phenotype in Ca_v_1.2^PFCKO^ mice is specific to social stimulus or generalizable to other rewards we tested mice using a non-social reward. The three-chamber test was replicated, but instead of a novel age-matched conspecific, the stimulus was a novel object that had been soaked in wet food pellets so that it emitted a potentially rewarding smell (**Figure 6A**). The opposite chamber contained a neutral-smelling novel object. The investigation in the first bout was similar across animal groups with mice spending more time in the zone around the food object (**Figure 6B**; *two-way* ANOVA, borderline main effect of zone F_1,22_=3.947, p=0.0596; no main effect of viral treatment, F_1,22_=0.6032, p=0.4456). During return investigations, both groups developed a significant preference for the food object (**Figure 6C**; *two-way* ANOVA, main effect of zone F_1,22_=26.18, p<0.0001; no main effect of virus F_1,22_=0.003453, p=0.9537). Analysis of the associated neural activity presented a more complicated picture. During the first bout, there was no significant difference in PFC activity in Ca_v_1.2^PFCGFP^ or Ca_v_1.2^PFCKO^ mice, either when actively interacting with the food stimulus versus object or not interacting (**Figure 6D**). During return bouts, there was a significant increase in PFC activity in Ca_v_1.2^PFCGFP^ mice during active interaction with the food stimulus compared to object that was absent in Ca_v_1.2^PFCKO^ mice (**Figure 6E, right**). No difference was observed while animals were not interacting (**Figure 6E, Left**). Quantification of the approach signal taken from −1s to +3s after entering the food zone revealed that both animal groups had a significant increase in PFC activity during the first bout (**Figure 6F**). However, during return bouts only Ca_v_1.2^PFCGFP^ mice had enhanced PFC activity upon entry into the food zone relative to neutral zone (**Figure 6G, left**). No difference was seen in Ca_v_1.2^PFCKO^ mice (**Figure 6G, right**). The lack of impairment in food reward approach behavior during return bouts (**Figure 6C**) compared to social preference behavior (**Figure 1E**) demonstrates that the type of reward stimulus is relevant, however the neural activity data suggests that despite the higher time spent with food stimulus in Ca_v_1.2 deficient mice there is a blunting of PFC activity.

**Figure 6.**
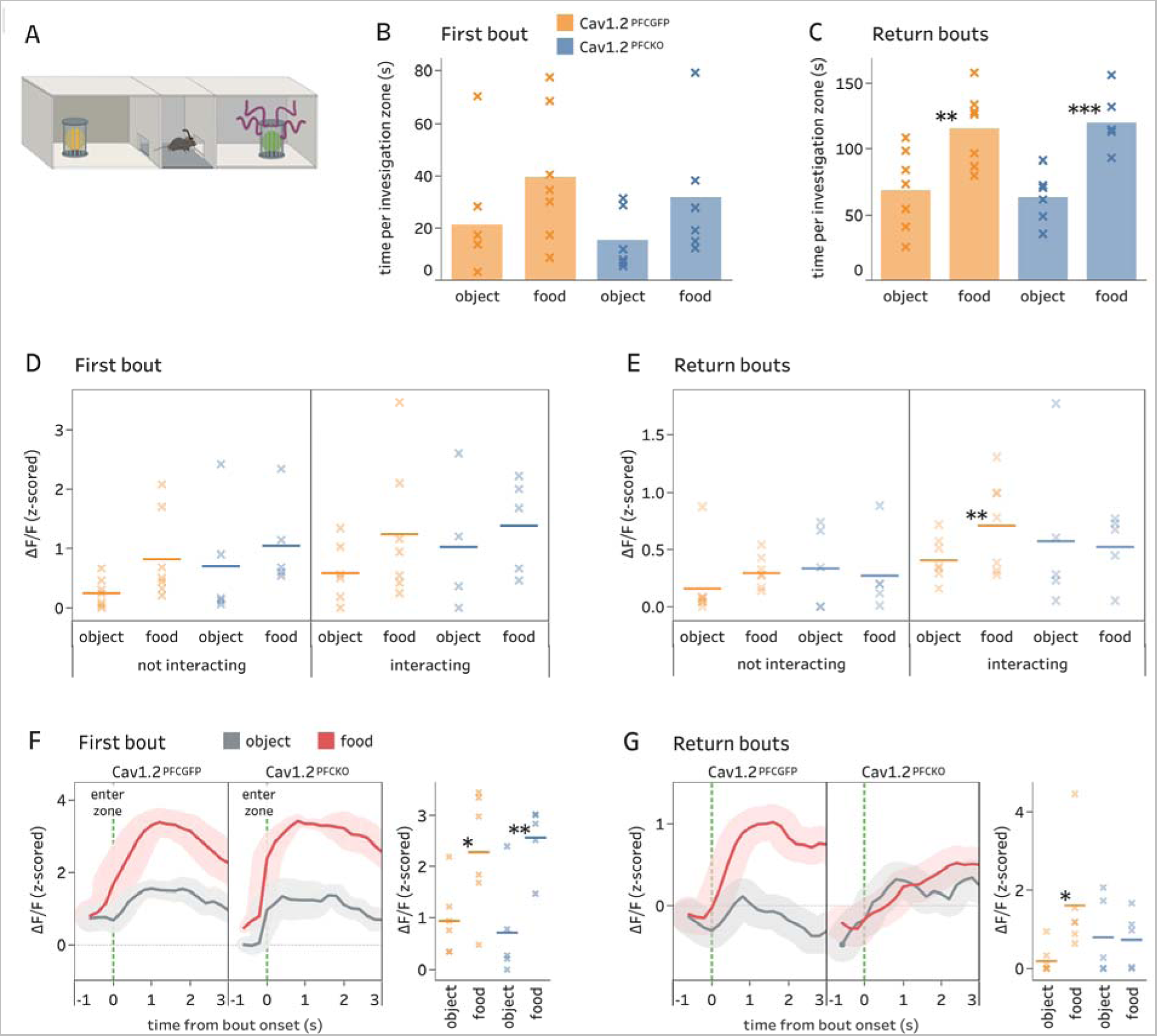
Food reward in the three-chamber approach test does not produce a behavioral deficit in Ca_v_1.2^PFCKO^ mice despite differences in neuronal activity. **6A.** Three-chamber food approach apparatus. **6B.** Food reward preference in the first bout, as measured by time investigating food-smelling object vs neutral novel object. Both animal groups spent more time with the food object in the first bout with no difference across genotype (Ca_v_1.2^PFCGFP^ n=7, Ca_v_1.2^PFCKO^ n=6). **6C.** For food reward preference in return bouts, there is no difference in time spent revisiting the food reward zone between Ca_v_1.2^PFCGFP^ and Ca_v_1.2^PFCKO^ mice (two-way ANOVA, genotype x investigation zone, P=0.61), with both showing a food preference (Bonferroni *post hoc* test, *P<0.05, **P<0.01 vs object; Ca_v_1.2^PFCGFP^ n=7, Ca_v_1.2^PFCKO^ n=6) **6D.** PFC GCaMP6s ΔF/F signal while interacting during the first bout does not vary across animal group or stimuli. **6E. Right**, In return bouts, ΔF/F while interacting is significantly higher when engaging with the food stimuli in Ca_v_1.2^PFCGFP^ mice but not Ca_v_1.2^PFCKO^ (Linear mixed effects model, Ca_v_1.2^PFCGFP^: **p = 0.00476, number of bouts = 45, n = 6; Cav1.2^PFCKO^: p = 0.4364, number of bouts = 37, n = 5). **Left**, No difference in ΔF/F when animals were not interacting (Ca_v_1.2^PFCGFP^: p = 0.1195, number of bouts = 53, n = 6; Cav1.2^PFCKO^: p = 0.5151, number of bouts = 39, n = 5). **6F. Left.** ΔF/F traces on entering investigation zones in first bout. **Right.** Quantitation of ΔF/F signal. Ca_v_1.2^PFCGFP^ and Ca_v_1.2^PFCKO^ mice both show a greater increase in entry signal on entering the food interaction zone for the first time relative to novel object (Linear mixed effects model, Ca_v_1.2^PFCGFP^: *p = 0.01152, number of events = 58, n = 6; Ca_v_1.2^PFCKO^: p = 0.00261, number of bouts = 66, n = 5). **6G. Left.** ΔF/F traces on entering zones in return bouts. **Right.** Ca_v_1.2^PFCGFP^ but not Ca_v_1.2^PFCKO^ mice show a significant increase in entry signal for the food reward relative to novel object in return (Linear mixed effects model, Ca_v_1.2^PFCGFP^: *p = 0.03245, number of events = 46, n = 6; Cav1.2^PFCKO^: p = 0.5950, number of events = 39, n = 5).

## Discussion

In the present study we demonstrate that focal knockdown of Ca_v_1.2 (*cacna1c*) reduced the tendency of male mice to investigate the location of a conspecific in the three-chamber social approach test, replicating our previous finding [8]. During the first investigation, PFC Ca_v_1.2 deficient mice (Ca_v_1.2^PFCKO^) interacted with the social partner at the same rate as control mice, displayed no social deficit, instigated a similar number of social investigation bouts, and had similar levels of PFC neural activity. Additionally, in a reciprocal social interaction assay, Ca_v_1.2 deficiency did not reduce the number of interactions or the associated neuronal activity. Therefore, the general capacity for social preference or exploration was not impaired. Rather, the average time per investigation in Ca_v_1.2^PFCKO^ mice when revisiting the social partner was similar to that when revisiting the non-social stimulus (object). This lack of social preference over time was coincident with a lack of increase in the population activity of PFC neurons during social interactions. This was in contrast to control mice spending more time with social versus nonsocial stimulus during repeat investigation with higher PFC activity while interacting with the social stimulus. A possible explanation for the observed mutant phenotype is that Ca_v_1.2^PFCKO^ mice derive less reward from repeat social interactions as delineated by the reduction in PFC activity, thereby precipitating an accelerated loss of interest in the social partner and an earlier exit from the investigation zone.

The investigation behavior of mice in the three-chamber task is likely rewarding given the development of a social preference when revisiting stimuli. A reward deficit in Ca_v_1.2^PFCKO^ mice is consistent with theories employed to explain common social deficits in neuropsychiatric disorders such as ASD, which posit that social stimuli are less rewarding in patients, resulting in a lack of interest to continually seek out social interaction [27, 28]. The *CACNA1C* risk variant rs1006737 has also been linked with defective reward responsiveness in humans [29] as well as abnormalities in PFC structure and function [30, 31]. In rodents, a recent paper demonstrated that *cacna1c* haploinsufficient rats emit fewer social vocalizations during playful social interactions, with the authors suggesting a potential role for social reward [10, 11, 13]. Consistent with our findings, there were no overt difference in the reciprocal interactions, but deficits were revealed during social approach. We additionally show that the behavioral deficit in Ca_v_1.2 PFC deficient mice was concurrent with an accelerated decline in neuronal population activity during repeat social interactions and a failure of approach dynamics to encode social preference. The specific impairments we observe in social approach behavior suggests that the neuropsychiatric-associated deficits brought about by PFC Ca_v_1.2 deficiency may be related to aberrant neural representation of social stimuli relevant to reward value processing. This is consistent with schizophrenia patients showing similar responses to social and non-social stimuli [32–35] and studies suggest that this is not due to an overall blunted response to nonsocial stimuli, but rather a specific social reward-processing deficit in neuropsychiatric conditions [36, 37].

As a focal point for the integration of internal motivational states and external sensory cues, the PFC is thought to regulate reward-seeking behavior by encoding context-dependent value computations [38]. In social contexts, a recent study found that manipulation of PFC neurons projecting to the nucleus accumbens (NAc) disrupted social-spatial learning in a modified three-chamber linear arena [22]. In this study we saw a social deficit in Ca_v_1.2^PFCKO^ mice in a similar social linear arena test, suggesting that the diminished PFC neuronal population signal during social interactions could be associated with a disruption in PFC-NAc reward pathways. However, neural correlates of value are near ubiquitous throughout the prefrontal cortex and other brain regions [39]. Attention, arousal, and motor preparation have all been demonstrated to correlate with the value of stimuli [40, 41]. Particularly as we were recording from a broad spectrum of neurons not associated with any one pathway, the reduction in signal during interaction behavior could be closer to the downstream expression of a reward deficit rather than a causal factor. Future studies will examine impact of Ca_v_1.2 deficiency in prefrontal circuits.

There is a possibility for a more comprehensive link between altered neural dynamics in Ca_v_1.2^PFCKO^ mice and reward-seeking behavior based on the correlation detected in control mice between PFC activity during initiation of a social encounter and the duration of social repeat investigation in the three-chamber task. That there was no such relationship in Ca_v_1.2^PFCKO^ mice is perhaps indicative of a failure of social cues to adequately translate to positive reinforcement. Mapping social approach over space by using a linear track, we also found that PFC population activity increased as animals moved towards a social partner, as characterized in previous studies [22]. This rise in the GCaMP6s signal may correspond to the accumulation of social information relevant to reward. In Ca_v_1.2^PFCGFP^ mice, the increase in activity relative to distance from the social target preceded that of Ca_v_1.2^PFCKO^ mice, where the signal increases at the point just prior to interaction. This discrepancy may be evidence of a Ca_v_1.2^PFCKO^-induced loss of an ‘anticipatory’ signal relevant to reward value representation. This possibility of reward-related dysfunction is further supported by the lack of any effect in Ca_v_1.2^PFCKO^ mice on repeat investigation of a novel non-social stimulus in the PFC-dependent forced alternation task and associated increase in PFC activity.

The PFC has long been implicated in the regulation of social behavior [14] but using a food reward instead of a social partner also preferentially recruited PFC neurons in control mice during return investigation, with a reduction seen in Ca_v_1.2^PFCKO^ mice. The fact that the difference in neuronal activity did not translate into a behavioral deficit may be because appetitive motivations such as hunger overrode reward pathways. Alternatively, the higher fluorescent signal in control mice during return bouts for the food stimulus could indicate that the PFC with normal expression of Ca_v_1.2 channels may still be performing a value computation based on incoming sensory information, but perhaps not relevant to reward processing. In this scenario, the PFC output related to social but not food stimuli would be given sufficient weight to influence reward-seeking behavior. Perhaps tellingly, the food object entry activity during return bouts was not predictive of reward-seeking behavior (**Supplementary Figure 2**) as it was with the social stimulus (**Figure 3E**).

If repeat social investigations in the three-chamber task are associated with reward-seeking, the lack of a Ca_v_1.2^PFCKO^ related difference in behavior or neural dynamics during first investigation may indicate there is no deficit in this initial exploration phase. However, the behavioral deficit in repeat investigations could originate from impaired social reward learning in the first bout. As mutations in the *CACNA1C* gene have been linked with problems in facial emotional recognition [7, 42], another function known to be carried out by the PFC [43, 44], any future reward-seeking deficit in PFC Ca_v_1.2 deficient mice could be connected to a deterioration in social recognition. In Ca_v_1.2^PFCKO^ mice there was a negative correlation between PFC activity during approach and time spent on investigation in the first encounter. A social recognition deficit could be expressed through requiring additional time to process and classify a novel mouse. Under this assumption that the first social approach activity is partially a neural correlate of social recognition, we can postulate a mechanism for a malfunction in reward processing. In the first phase the animal would be acquiring a representation of social reward value, a process that is impeded in Ca_v_1.2^PFCKO^ mice. This initial inability to form a reward association distorts the PFC reward pathway and precipitates an accelerated decrease in response to rewarding stimuli during future social interactions and ultimately the observed deficit in social reward-seeking behavior [45, 46].

A model of impaired sociability observed in Ca_v_1.2^PFCKO^ mice that pertains to either reward responsiveness or social recognition deficit would be consistent with our current knowledge of neuropsychiatric disorders such as autism and schizophrenia that needs to be further explored. Here we have shown that a reduction in Ca_v_1.2 channels in the prefrontal cortex suppresses the development of a sustained social preference in mice. There are concurrent changes in neuronal population dynamics during social interaction and approach that may be relevant to reward-seeking behavior. To further elucidate the nature of this social deficit and how it relates to neuropsychiatric disorders requires identification of the specific pathways and cell types involved, as well as manipulation of neuronal populations to establish a causal link between Ca_v_1.2 channels and social approach behavior.

## Supplementary Figures

**Supplementary Figure 1.**
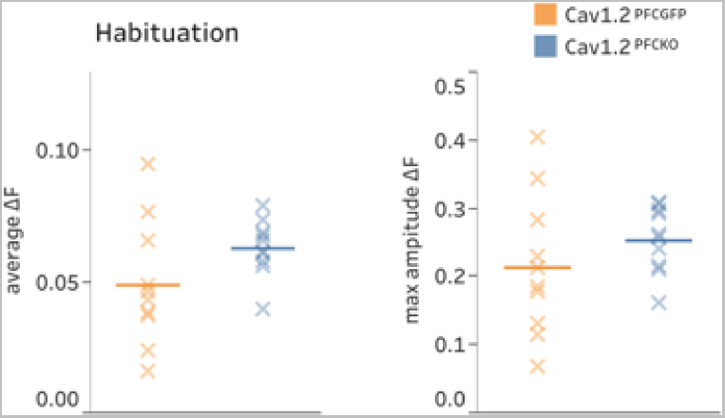
During the habituation period there was no difference in average PFC GCaMP6s fluorescence or amplitude between Cav1.2^PFCGFP^ and Cav1.2^PFCKO^ mice demonstrating that loss of Ca_v_1.2 channels did not impact GCaMP6s fluorescence.

**Supplementary Figure 2.**
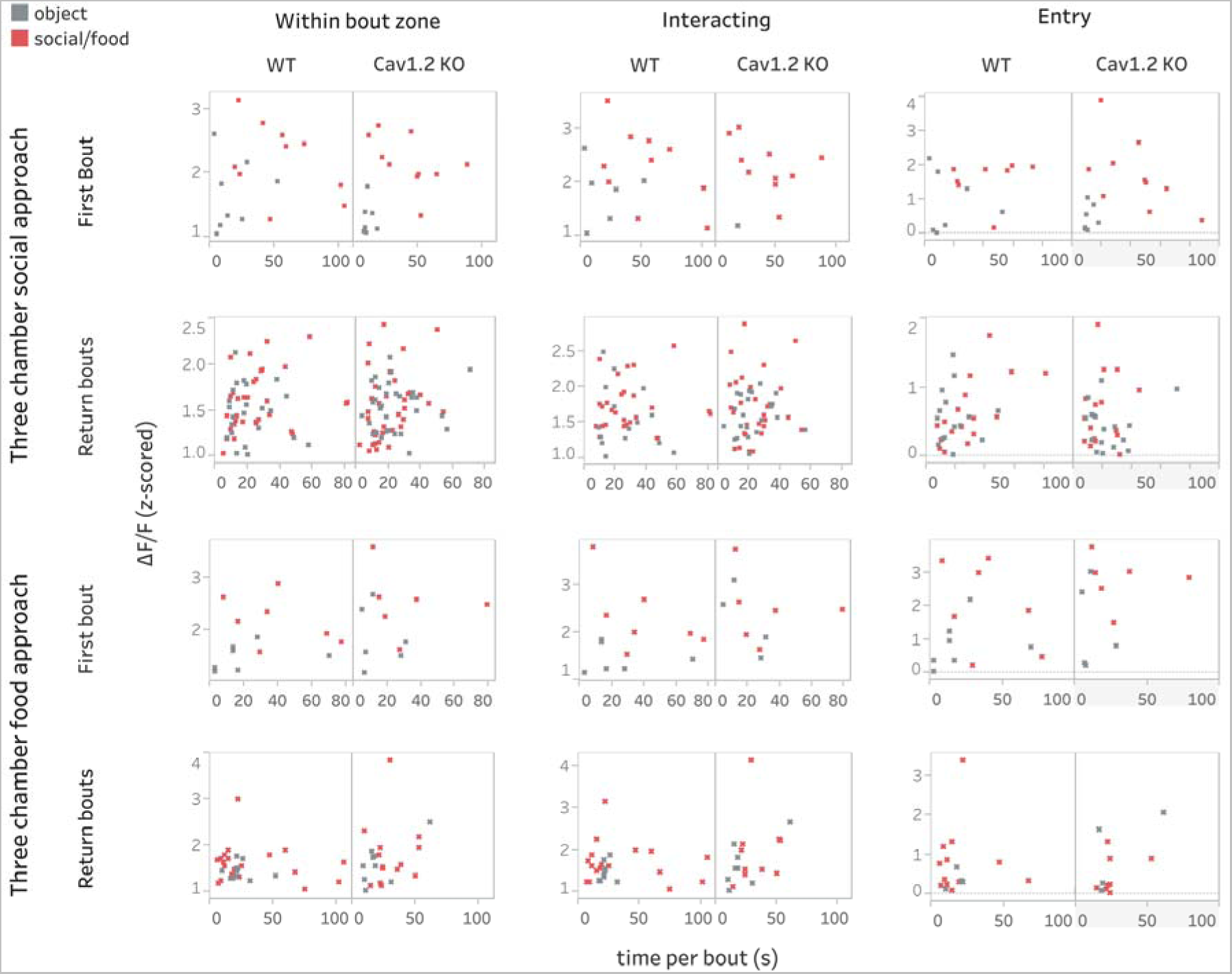
Correlation plot of within bout, interacting and entry signal during first and return bouts for three chamber social approach (top two panels) and three chamber food approach (bottom two panels) tests.

## Data and materials availability

All data are available in the main text or the supplementary materials. MATLAB code for fiber photometry data analysis is available upon request.

